# Structure of the CABIT2 domain of THEMIS reveals a novel protein fold with an inserted SH3-like domain

**DOI:** 10.1101/2024.10.01.616165

**Authors:** Tyler S. Beyett, Zeynep Yurtsever, Ilse K. Schaeffner, Jaimin K. Rana, Lucy K. Robbie, Daniela Ortiz-Salazar, Michael J. Eck

## Abstract

Maturation of thymocytes into T cells is critical for proper function of the adaptive immune system. During this developmental process, thymocytes undergo a highly-regulated selection process regulated by the signaling characteristics of the T cell receptor (TCR) pathway. Thymocyte-Expressed Molecule Involved in Selection (THEMIS) is an essential protein for T cell development. THEMIS regulates phosphatases downstream of the T cell receptor to ensure signaling thresholds are met during selection. Important features of THEMIS are its two uncharacterized CABIT (Cysteine-containing All-Beta In THEMIS) domains, which are intriguing because they have been proposed to participate in important protein-protein interactions (PPIs) that modulate immunological signals. Here, we report the 2.9 Å crystal structure of the THEMIS CABIT2 domain determined via heavy atom phasing. The structure revealed a novel protein domain fold comprised mainly of β-sheets with two distinct subdomains. This domain appears to have a different C-terminal boundary than predicted or found in previously used experimental constructs. Inclusion of the proline rich segment enables GRB2 to bind CABIT2. Isolated CABIT2 domain is unable to bind or modulate the function of SHP1 phosphatase. This structure will provide the foundation for future structure-function studies of CABIT domains and THEMIS.

## Introduction

T cells play an essential role in the adaptive immune system, recognizing antigens and carrying long term memory of previously encountered pathogens. Development of mature T cells from their immature thymocyte precursors occurs in the thymus and depends on the meticulous balance of cellular signaling of the T cell receptor (TCR) pathway.^1,2^ In the thymus, developing T cells first undergo positive selection to select for double positive (DP) thymocytes expressing both the TCR coreceptors CD4 and CD8. Cells subsequently undergo negative selection and mature into single positive (SP) thymocytes expressing only one of the coreceptors based on their binding affinity to self-peptide bound major histocompatibility complex (pMHC) and the resulting level of downstream intracellular TCR signal intensity.^2,3^ The complexity of this selection process necessitates tightly regulated TCR signaling to ensure sufficient receptor-antigen affinity to recognize pathogens but not lead to autoimmune diseases.^2,4^ This is followed by the final negative selection stage in which thymocytes are further differentiated into either cytotoxic or helper T cells.^1^

Themis (THymocyte-Expressed Molecule Involved in Selection) was discovered in a search for genes implicated in T cell development; its inactivation/knockout halted development at the positive selection stage leading to cell death.^5–9^ THEMIS was found to be a crucial regulator of signaling downstream of the TCR and thymocyte maturation, especially at the positive selection stage where its protein levels are highest.^10^ A B cell-specific isoform THEMIS2 has been identified that regulates B cell receptor (BCR) signaling.^11–14^ Development of THEMIS^− /–^ T cells could can be recovered by the expression of THEMIS2, suggesting it has a similar functional role in regulating TCR and BCR signaling.^14^

Co-immunoprecipitation experiments showed THEMIS is frequently found in complex with SHP1 phosphatase and the adaptor protein GRB2, though whether GRB2 is required for THEMIS to interact with SHP1 is debated.^15–19^ Current models of TCR regulation posit that GRB2 interacts simultaneously with THEMIS, SHP1, and the phosphorylated linker for activation of T cells (LAT) to localize the complex to the membrane and in proximity of TCRs.^7,15,17^ However, the models diverge and offer competing and contradictory claims about how THEMIS modulates TCR signaling.^10^ The first model argues that THEMIS activates SHP1 to decrease signaling strength through the TCR.^5,17,20–23^ Recent evidence showed that lymphocyte-specific protein tyrosine kinase (LCK) can phosphorylate THEMIS that can act as a “priming substrate” to activate SHP1.^23^ The second model argues that THEMIS inhibits SHP1 to enhance TCR signaling by catalyzing the oxidation of the active site cysteine in SHP1 required for catalysis.^16,18,19,24^ Reactive oxygen species (ROS) are generated during TCR signaling and may serve as the oxidizer in this model.^25^ In either case, the resulting TCR signaling strength is sufficient for positive selection and proper maturation, suggesting the affinity of the pMHC ligands could determine whether TCR signal dampening or enhancement occurs.

A major challenge associated with the study of the function of THEMIS is the lack of structural or biochemical characterization, as nearly all prior work has been cell based. THEMIS is almost entirely composed of two Cysteine-containing, All-Beta-In-THEMIS (CABIT) domains that are predicted to only be present in one other protein family, GRB2-associated and regulator of Erk/MAPK (GAREM), of which there are two isoforms.^10,26^ GAREM regulates the MAP kinase/Erk signaling pathway also through interactions with SHP phosphatases.^27^ Structural predictions of the CABIT domains of THEMIS suggest this uncharacterized domain is comprised of nearly all beta sheets and does not resemble any reported protein fold. The function of CABIT domains is also unknown, but they are not predicted to have enzymatic activity. Rather, they likely modulate signaling through protein-protein interactions (PPIs) with other enzymes like SHP1. The N-terminal CABIT domain (CABIT1) has been shown to interact with SHP1, and a proline-rich segment (PRS) immediately following the second CABIT domain (CABIT2) is the binding site of GRB2.^15,17,23,24^ The C-terminal tail following the PRS spans approximately 100 residues and is predicted to be disordered.

To better understand the function of THEMIS and begin to clarify how it regulates TCR signaling, we determined the 2.9 Å crystal structure of the CABIT2 domain of THEMIS phased using heavy atom soaking. This represents the first experimental structure of a CABIT domain, which revealed a novel domain fold. We show that previously predicted CABIT2 domain boundaries and some reported experimental constructs omit critical secondary structure features, likely disrupting folding of the CABIT domain. We report alternative constructs that are folded and biochemically well-behaved. We show the CABIT2 domain containing the PRS interacts with GRB2 but not SHP1, and the CABIT2 domain has no effect on the enzymatic activity of SHP1. The constructs and crystallization methods we report here will facilitate future studies on this novel domain and aid in resolving longstanding questions in the THEMIS field.

## Results

### Defining the domain boundaries of the CABIT2 domain

While the N-terminus of the human CABIT2 domain is commonly reported as residue 267, the C-terminal boundary is not agreed upon. Predictions by Hidden Markov modeling and Uniprot describe the terminus as residues 529 or 518, respectively (Figure 1A). However, constructs spanning 267-530 and 267-520 are insoluble when expressed in *E. coli* likely due to the removal of secondary structure features important for domain folding (Figure 1A, B; SI Figure 1A). Furthermore, a construct spanning residues 1-493 has been used in cellular and *in vivo* studies to represent CABIT1+CABIT2 (Figure 1 A, B), but our attempts to purify a construct spanning 267-493 yielded insoluble protein (SI Figure 1A).^19,23,28^ While many of these reported constructs utilize mouse THEMIS, it is predicted to have the same domain topology and residue positioning as human THEMIS. Using secondary structure predictions to create alternative constructs, we generated a construct spanning 267-550 that can be readily purified from bacteria and is thermally stable (T_m_=51.34 ± 0.03 °C, SI Figure 1A, B; SI Figure 1).

**Figure 1:**
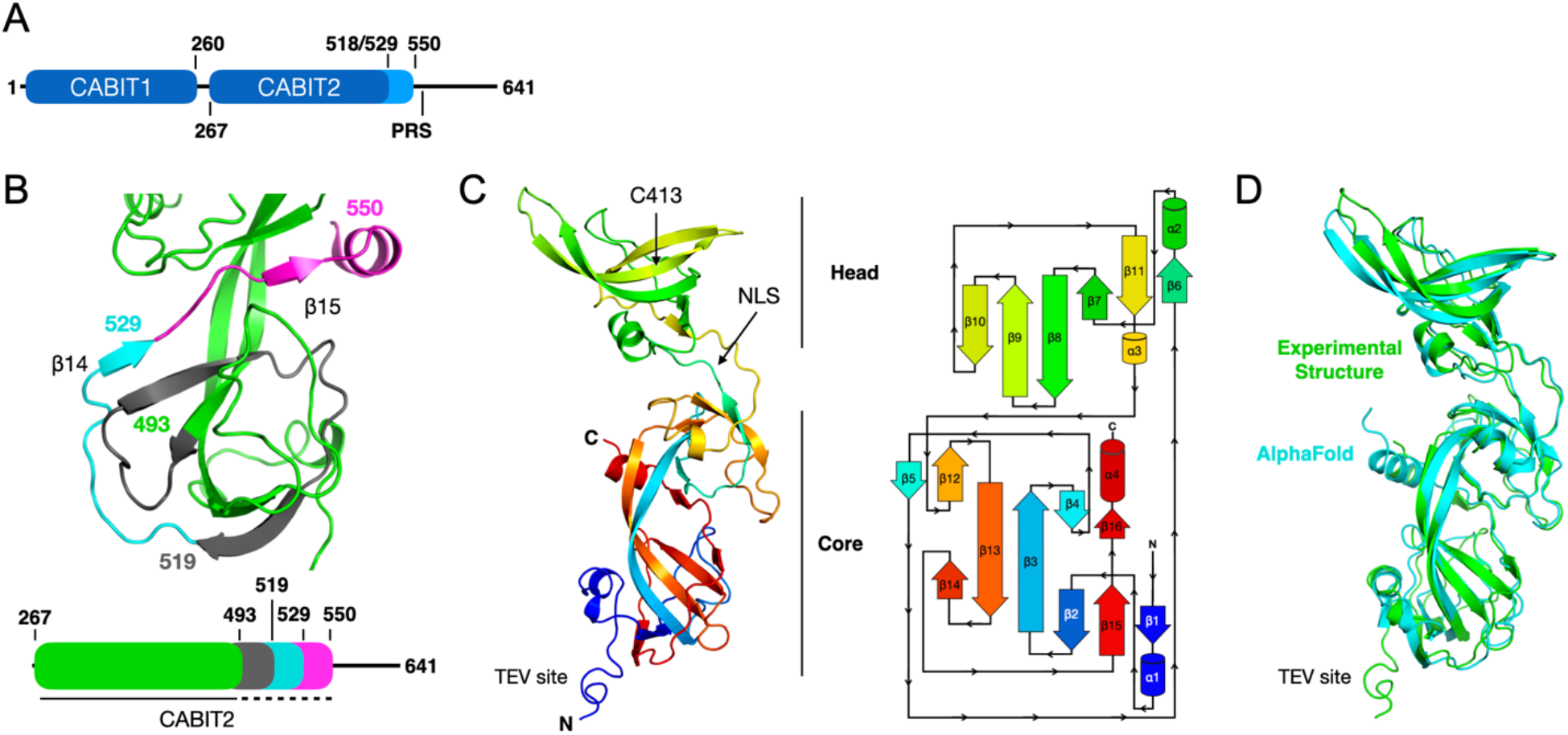
Crystal structure of the CABIT2 domain. A) Domain layout of THEMIS. The construct crystallized spanned residues 267-550, which is longer than predicted (518 and 529) for the CABIT2 domain. Following CABIT2 is a proline rich segment (PRS) ending ∼564 that GRB2 binds. B) Different C-terminal truncations for the CABIT2 domain are shown on our CABIT crystal structure. The crystallized construct truncating at residue 550 encompasses the entire structure shown. Truncation at 529 deletes the magenta segment, truncation at 519 deletes the magenta and cyan segments, and truncation at 493 deletes the magenta, cyan, and gray segments. In each case, truncation removes secondary structure elements and leads to insoluble protein. C) Gold SAD phased CABIT2 structure at 2.9 Å. The protein is colored in rainbow from N-terminus (blue) to C-terminus (red). The core and SH3-like head subdomains are indicated, as are conserved C413 and a putative NLS. D) Global superposition in PyMol of the AlphaFold predicted CABIT2 domain structure (residues 267-550, cyan) and experimentally determined structure (green). The cyan tail without an aligned prediction is the TEV site from the crystallized construct.

### Crystallization and structure determination

Given our inability to purify soluble protein using predicted boundaries of the CABIT2 domain, we opted to use our construct spanning residues 267-550 for crystallization due to its solubility and expression levels. Sparse-matrix screening with purified CABIT2 yielded >50 conditions that produced crystals with a hexagonal cylinder morphology (SI Figure 2A) but only with an attached N-terminal strep tag and TEV protease cleavage site. After optimization of several conditions, we obtained the best diffracting crystals in 0.1 M imidazole pH 6.9 and 1.0 M sodium acetate. Large crystals (>400 µm in length) diffracted to ∼3 Å, but we were unable to determine the structure via molecular replacement through an automated search using the entire PDB. As these diffraction data predated AlphaFold, we attempted to phase the data using structure predictions from the PHYRE and iTASSER servers but were unsuccessful.^29,30^ Thus, we experimentally phased the structure using single-wavelength anomalous diffraction (SAD) with selenomethionine incorporation or heavy atom soaking with gold or mercury complexes. Here, we report the gold-phased structure of the THEMIS CABIT2 domain to 2.9 Å resolution due to its superior electron density quality and higher resolution compared to other SAD phasing results (Figure 1C). After successfully determining the structure, a prediction of the CABIT2 domain from AlphaFold was released that allowed for phasing to be accomplished via molecular replacement (Figure 1D).^31^

SAD phasing of the CABIT2 domain was complicated by weak anomalous signal and a long unit cell axis (>400 Å) that required careful detector placement to avoid spot overlap (SI Figure 2B). We collected and merged three data sets from a single crystal soaked in K[Au(CN)2] to obtain a highly redundant data set, aided by the high symmetry P6_1_22 space group, with sufficient anomalous signal for phasing in Phenix with AutoSol (SI Table 1, SI Figure 2A). Using the density-modified map generated from SAD phasing (SI Figure 2C), an initial model was prepared with Phenix AutoBuild that was manually rebuilt and refined to obtain the final structure (Figure 1C, SI Table 1, SI Figure 2D). Strong positive density corresponding to gold is observed near coordinating Cys and His residues (SI Figure 2D). There is one molecule of the CABIT2 domain per asymmetric unit (ASU), which leads to an unusually high solvent content of 75% and likely contributes to the weak anomalous signal. The overall CABIT2 domain architecture we experimentally determined aligns well with the predicted structure from AlphaFold with an overall root mean square deviation (RMSD) of 1.7 Å (Figure 1D). Most differences between the predicted and experimentally determined structures are in regions involved in crystal contacts.

### Features of the CABIT2 domain structure

Despite its name, the CABIT domain is not all-beta but is nearly all beta, with 16 β-sheets and four short helices (Figure 1C). The CABIT2 domain has two distinct subdomains. The “core” subdomain is comprised of the N- and C-terminal regions (approximately residues 267-344 and 458-544, respectively) of the polypeptide chain and resembles a pair of small β-barrels that share two central extended antiparallel β-sheets (β3 and β13). We suspect that the core being composed of discontinuous regions of the polypeptide chain may have been responsible for poor structure predictions prior to AlphaFold. The “head” subdomain spans approximately residues 345-457 and resembles an SH3 domain with an unusually long pair of antiparallel sheets (β8-β9). The universally conserved cysteine residue for which the CABIT domain is named (Cys 413 in the CABIT2 domain) is buried in the hydrophobic core of this SH3-like subdomain (Figure 1C). Thus, it may be conserved simply for structural reasons – to preserve the hydrophobic packing required for the proper folding of this subdomain.

The 7-residue TEV protease site remaining on our uncleaved CABIT2 domain construct forms a short helix-like segment at the N-terminus of the core subdomain (Figure 1B). Inspection of the crystal lattice reveals that this segment is involved in crystal packing interactions with two adjacent CABIT2 domains (Figure 2A, SI Figures 2C, E). The participation of the TEV site in crystal contacts explains our inability to obtain crystals after TEV protease cleavage to remove the N-terminal strep tag and this protease recognition sequence.

**Figure 2:**
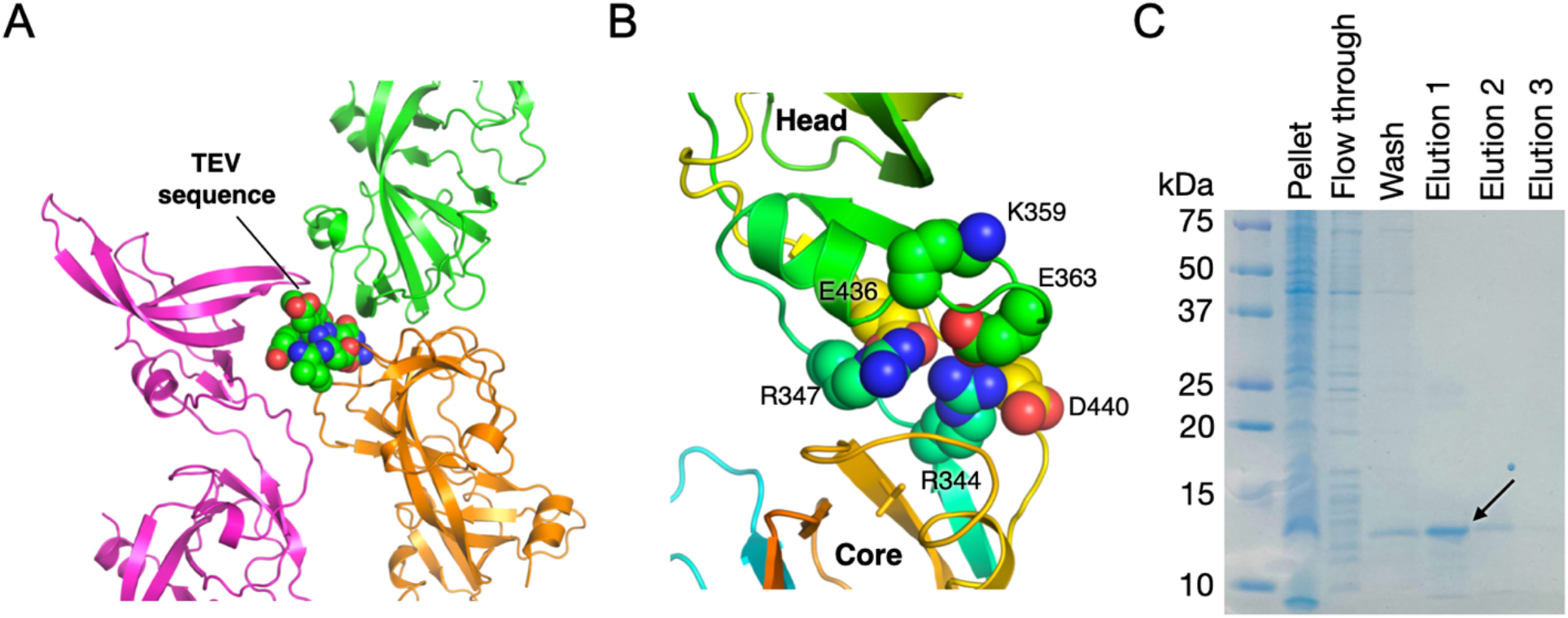
Features of the CABIT2 structure. A) Close-up view of the crystal contact involving the TEV sequence (shown in spheres). This region assumes a poorly ordered helix and contacts two adjacent protomers. B) The interface between the core and SH3-like subdomains. Numerous charged residues are found in the core. C) The SH3-like head subdomain (residues 347-436) can be purified as an isolated and soluble protein.

Themis has been predicted to contain a nuclear localization signal (NLS) that has been proposed to be monopartite (residues 345-349) or bipartite (residues 330-332 and 345-349) in nature.^14,32^ The present structure shows that this highly basic segment lies at the interface of the core and SH3-like subdomains and forms the first β-strand of the latter and is therefore unlikely to be sufficiently exposed to function as an NLS, at least in the conformation observed in our crystal structure (Figure 1C). R347 in this segment forms a salt bridge with E436 and R344 in the core subdomain forms a salt bridge with E363 in the SH3-like domain. These charged residues, along with D440, Y442 and K469, form most of the interface between the core and SH3-like domains. There are few hydrophobic contacts in this interface, suggesting to us that the SH3-like could be an independently folded domain (Figure 2B). Consistent with this notion, we found that a construct spanning residues 347-436 was readily purified and soluble (Figure 2C).

### CABIT2 domain conservation and homology

The overall fold of the CABIT2 domain of THEMIS is unusual and prompted us to perform a structural similarity search with the DALI server.^33^ While portions of the CABIT2 domain have weak homology to portions of other protein structures, we were unable to identify any published structures with homology to the entire CABIT2 domain leading us to label it as a novel fold. Since THEMIS is conserved across metazoans, we used the CONSURF server to identify regions of the protein with evolutionary conservation that can, for the first time, be accurately mapped onto a structure (Figure 3A, SI Figure 3).^34,35^ The hairpin loops found in the SH3-like subdomain are poorly conserved, as are two loops in the portion of the core formed by the N- and C-termini (residues 290-300 and 515-530, respectively). The most highly conserved residues are located along the interface between the core and SH3-like subdomains, the putative NLS, and C-terminal helix.

**Figure 3:**
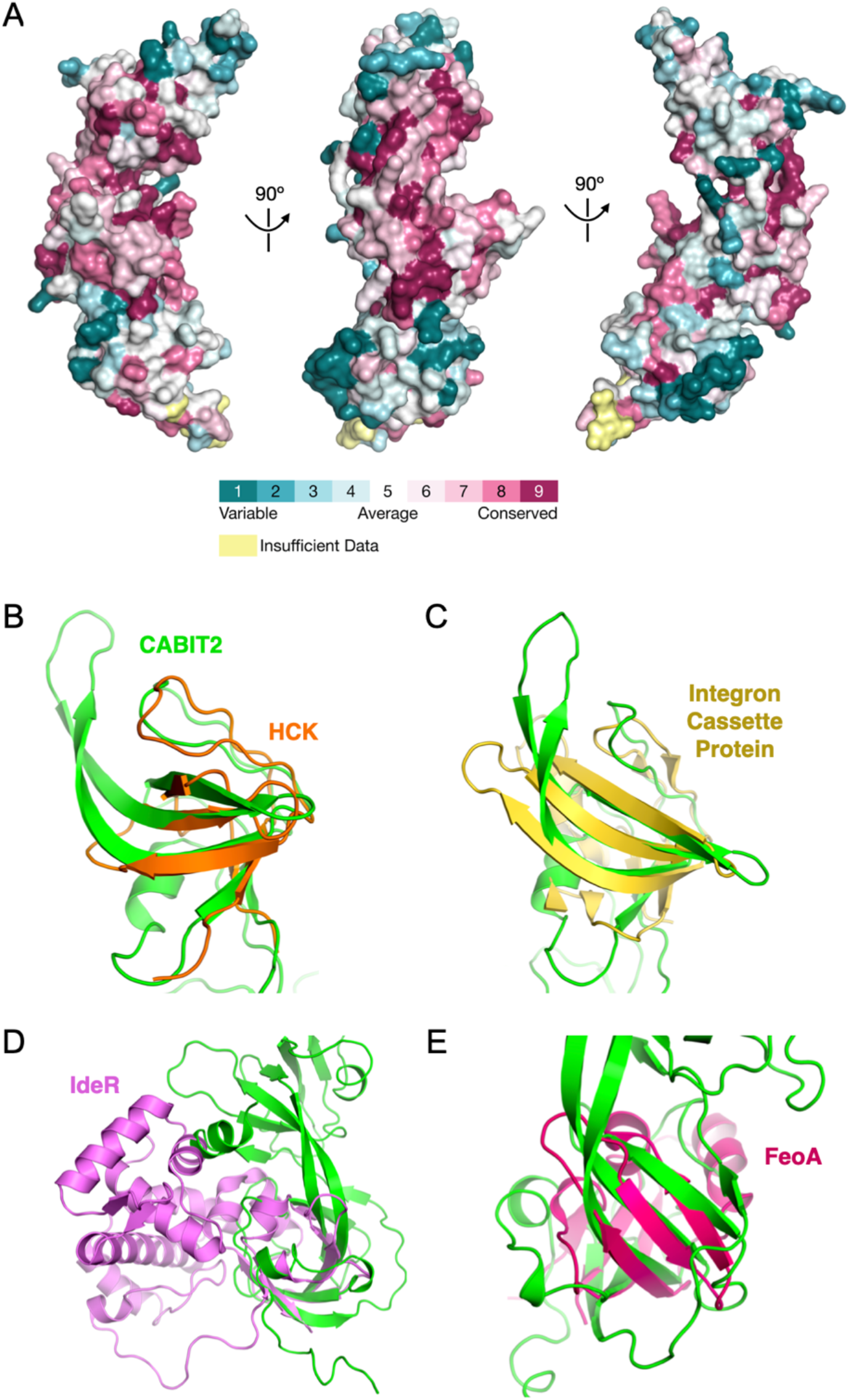
Conservation of the CABIT2 domain. A) Surface representation of the CABIT2 domain colored according to sequence conservation as determined by the CONSURF server. Insufficient data corresponds to much of the synthetic TEV site. B) Structural alignment between the SH3-like head subdomain of CABIT2 and the SH3 domain of HCK (PDB 3REA, RMSD=2.8 Å). Structural conservation was determined by the DALI server. C) Structural alignment between the SH3-like head subdomain of CABIT2 and a *Vibrio cholere* integron cassette protein (PDB 3G1J, RMSD=2.8 Å). D) Structural alignment of IdeR and the core of the CABIT2 domain (PDB 1FX7, RMSD=2.5 Å for overlapping region only). E) Structural alignment between FeoA and the core of the CABIT2 domain (PDB 3E19, RMSD=2.5 Å).

DALI structural similarity search results revealed the SH3-like head indeed aligns with many SH3 domains from proteins including LCK, FYN, SRC, and HCK (Figure 3B). However, the extended hairpin between β8-β9 is not found in these canonical SH3 domains and has only been observed in the structure of an integron cassette protein from *Vibrio cholerae* (Figure 3C). Several loops and the α2 helix of the head would interfere with canonical SH3 substrate binding, suggesting that this subdomain may not bind polyproline peptide substrates. The β-barrel like portion of the CABIT2 domain core near the N- and C-termini has weak structural homology to several metal-dependent transcriptional regulators of the diphtheria toxin repressor (DtxR) family including IdeR, MtxR, ScaR, and SloR (Figure 3D) and components of the bacterial ferrous iron transport system like FeoA (Figure 3E). Despite these weak structural similarities, there is no evidence that CABIT domains bind endogenous metals.

### Insights into the structure of CABIT domains from AlphaFold

Surface electrostatics analysis revealed one face of the core to be highly acidic (Figure 4A). Much of this surface is predicted with high confidence by AlphaFold to be in contact with the THEMIS CABIT1 domain (Figure 4B). A basic patch predicted to be an NLS is located on the other side of interface between the core and SH3-like subdomains including residues K343, R345, K441, and K460. The loop connecting β3-β4 (residues 323-329) is unusual, as it projects F327 in a strained rotameric conformation into solvent. In the predicted full-length structure of THEMIS, this exposed loop is predicted to be stabilized by a cation-π or backbone hydrogen bond with K208 in the CABIT1 domain (Figure 4C).

**Figure 4:**
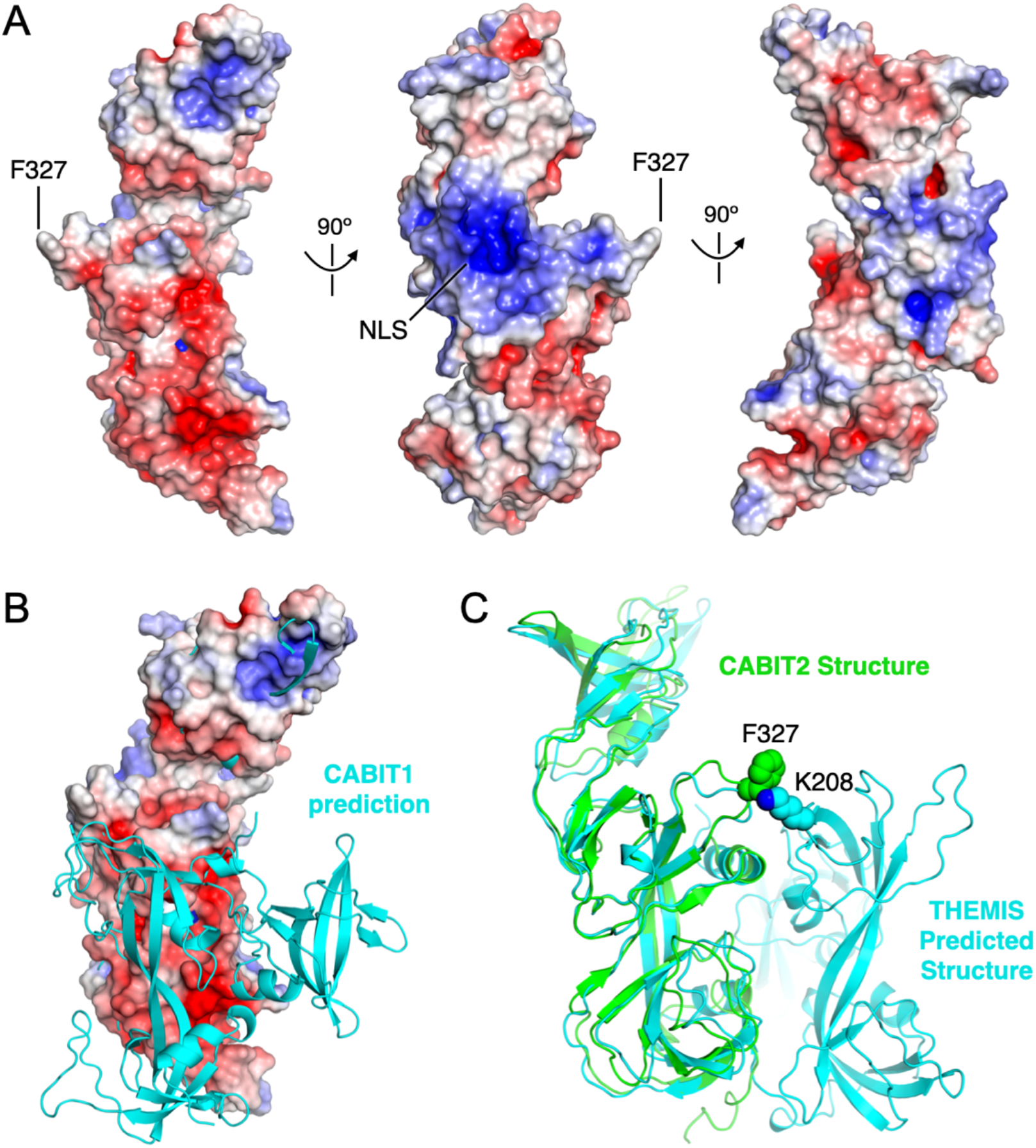
Electrostatics of the CABIT2 domain. A) Electrostatic surface of the CABIT2 domain. Positive and negative charges are blue and red, respectively. B) The predicted location of the CABIT1 domain in THEMIS is near the acidic patch of CABIT2. C) F327 is extended into solvent in the isolated CABIT2 domain but is predicted to make favorable contacts with K208 in the CABIT1 domain.

THEMIS is predicted to have two CABIT domains. Comparison of our CABIT2 crystal structure with the AlphaFold CABIT1 prediction revealed the same general organization with a core featuring a pair of extended antiparallel β-sheets and an SH3-like subdomain (Figure 4B; SI Figure 4A). Unlike the CABIT2 domain structure in which the SH3-like subdomain is in line with the core, the CABIT1 SH3-like subdomain is predicted to be oriented perpendicular to the core and lacks the extended β-hairpin found in the CABIT2 domain. The rotation of the central β-sheets in the predicted CABIT1 domain core is opposite of the rotation in our CABIT2 structure.

The CABIT2 domain in THEMIS2 and THEMIS3, the latter of which is not found in primates, are predicted to closely resemble that of THEMIS with an expected RMSD of 3.3 Å and 3.6 Å, respectively (SI Figure 4B, C). GAREM, of which there are two isoforms, is the only other human protein family predicted to contain a CABIT domain. Hidden Markov modeling predicts the GAREM CABIT domain to span residues 1-344, which is approximately 60 residues longer than the crystallized CABIT2 domain in THEMIS. The predicted fold of the GAREM CABIT domain includes an approximately 30 residue insertion into the extended β8-β9 hairpin observed in the SH3-like subdomain of THEMIS CABIT2 in addition to an additional antiparallel β-sheet (SI Figure 4D). The predicted core of GAREM more closely resembles that of the CABIT2 domain than the CABIT1 domain in THEMIS, but there are still notable differences in the predicted threading, especially near the C-terminus of the CABIT domain where there are helices that account for approximately 30 additional residues compared to THEMIS CABIT2. Thus, while predictions suggest THEMIS and GAREM CABIT domains have conserved subdomains, there is likely enough structural variability to allow for a variety of functions and protein-protein interaction.

### The CABIT2 domain does not bind or activate SHP1

CABIT domains are not known to have any intrinsic enzymatic activity. Rather, THEMIS is proposed to modulate the activity of SHP family phosphatases through protein-protein interactions with its CABIT domains. Accordingly, we assessed the activity of SHP1 phosphatase in the presence of varying concentrations of purified CABIT2. The THEMIS CABIT2 domain alone did not exhibit measurable phosphatase activity (Figure 5A), nor did it affect the activity of full-length SHP1 (Figure 5B). However, it unexpectedly enhanced the activity of the truncated SHP1 PTP domain construct (Figure 5C). A lack of an effect on full-length SHP1 was expected, as a recent report showed phosphorylation of Y34 in the CABIT1 domain is important for modulation of SHP1 activity in an allosteric manner.^23^ The mechanistic basis for the effect with observe on the isolated phosphatase domains of SHP1 is unclear.

**Figure 5:**
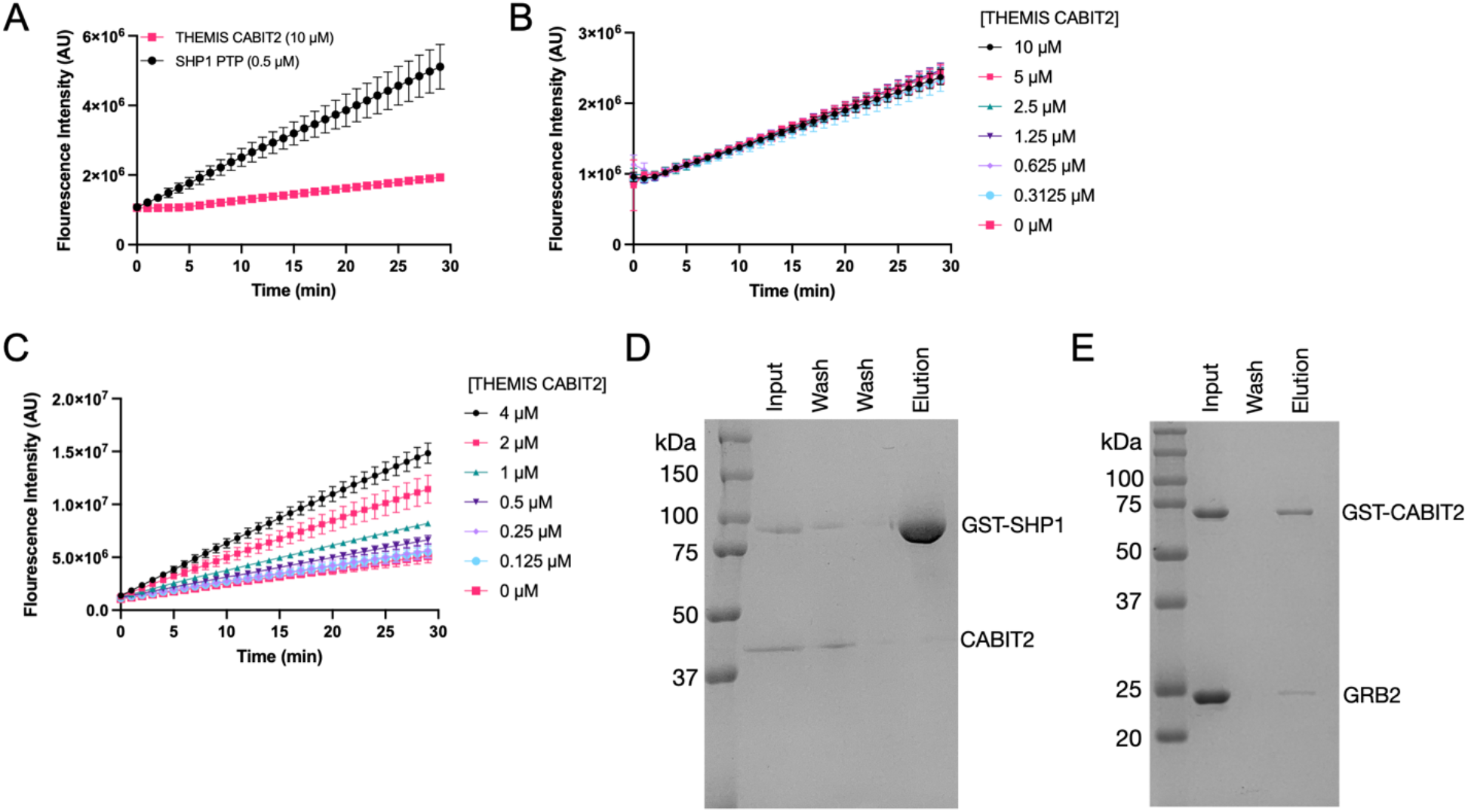
Functional and interaction assays with isolated THEMIS CABIT2 domain. A) Phosphatase assay using the fluorogenic substrate DiFMUP, whose fluorescence increases upon dephosphorylation. THEMIS CABIT2 domain does not display phosphatase activity. The PTP domain of SHP1 was used as a positive control (n=3, reported as mean±SD). B) Titration of THEMIS CABIT2 in the presence of a constant concentration of full-length SHP1 (3.5 µM) does not affect the activity of SHP1 (n=3, reported as mean±SD). C) Titration of THEMIS CABIT2 in the presence of a constant concentration of SHP1 PTP domain (0.5 µM) increases enzymatic activity (n=3, reported as mean±SD). D) Pulldown with GST-tagged SHP1 and THEMIS CABIT2 domain. CABIT2 and SHP1 do not interact. E) Pulldown with GST-tagged THEMIS CABIT2 containing the PRS (267-564) and GRB2. A weak interaction is observed.

We next assessed the ability of the CABIT2 domain to bind to SHP1 and GRB2, the most studied binding partners of THEMIS. We used a slightly longer construct (residues 267-564) containing the proline-rich segment (PRS) for these experiments, as the PRS has been shown to be required for GRB2 association.^17^ In pulldowns using GST-tagged CABIT2, no association between THEMIS and full-length SHP1 was observed (Figure 5D). Weak association with GRB2 was observed (Figure 5E). Together, these results show that the CABIT2 domain containing the PRS weakly interacts with GRB2 but does not interact with or affect the enzymatic of full-length SHP1.

## Discussion

THEMIS represents one of four known proteins in humans to contain a CABIT domain, the others being THEMIS2, GAREM, and GAREM2, and all are known to modulate tyrosine kinase signaling through PPIs with SHP family phosphatases and GRB2. THEMIS and THEMIS2 are expressed in T and B cells, respectively. GAREM (FAM59A) is ubiquitously expressed and GAREM2 (FAM59B) is neuronally restricted.^26,27^ While the precise molecular mechanisms underlying signal modulation remain unclear, the novel fold we reveal here suggests modulation and PPIs may occur through novel mechanisms. Thus, CABIT domains represent important new folds to study with high relevance to adaptive immunology and beyond.

Our THEMIS CABIT2 structure was determined using experimental phasing and is highly similar to the AlphaFold prediction for this same construct (RMSD=1.7 Å) including most residue side chain positions. The main differences were in regions participating in crystal contacts. This impressive result instills confidence in AlphaFold predictions of other CABIT domains. However, it remains to be seen how CABIT domain tandems in THEMIS and THEMIS2 are arranged. The prediction for THEMIS has both CABIT domains in contact with one another, whereas the prediction for THEMIS2 has them not in contact.

Our CABIT2 structure revealed a novel fold with two distinct regions that we have termed the “core” and the “head”. The head subdomain is SH3-like in structure as predicted by the DALI server but does not have full homology to any reported SH3 domains. The canonical polyproline substrate binding site is occluded by a helix indicating it must have another function. The SH3-like head subdomain appears to be inserted into the core. Indeed, the head subdomain can be excised from the CABIT2 domain and purified as a soluble protein. We speculate that the head subdomain may have been the result of a genetic insertion event of an SH3-like domain into the core. A similar event has been observed in G protein-coupled receptor (GPCR) kinases (GRKs), which have an AGC family kinase domain inserted into a loop of a regulator of G protein signaling homology (RH) domain.^36,37^

A notable result of our structural work concerns the domain boundaries of the CABIT2 domain. The present structure shows that the C-terminal boundary of the CABIT2 domain is approximately residues 550. We were unable to purify shorter constructs spanning the previously predicted domain range (residues 267-530 or 267-520) and the structure indicates that these constructs lack secondary structure features integral to the fold of the core of the CABIT2 domain. We note that several prior studies of THEMIS function have employed a construct spanning residues 1-493 for the CABIT1-CABIT2 domain tandem^19,23,28^ and one has used 521 as a C-terminal cutoff for CABIT2.^32^ Our attempts to purify CABIT2 using residue 493 or 520 as the C-terminal cutoff yielded insoluble protein. These constructs lack several secondary structure features in the core subdomain that likely impact domain folding. It is possible that a construct spanning 1-493 will contain a functional CABIT1 domain, but our results suggest that the CABIT2 domain in such a construct will be unfolded. Additionally, this construct does not span the CABIT2 domain regardless of whether the predicted (residue 529 or 518) or crystallized (residue 550) positions are considered as the domain boundary. Thus, prior cellular and *in vivo* work with these shorter constructs should be re-evaluated with this caveat in mind.

How THEMIS regulates the activity of SHP phosphatases is debated in the literature.^10^ THEMIS may both activate and inhibit SHP1 under different physiological conditions. Our phosphatase assays with purified CABIT2 domain revealed that this domain does not affect the activity of full-length SHP1. This is expected based on recent literature showing that CABIT1 interacts with SHP1.^23^ To our surprise, we found that CABIT2 enhanced the activity of the isolated phosphatase domain of SHP1, which is already highly active due to truncation of the regulatory SH2 domains. The physiologic relevance of this observation is unclear. While the truncated PTPase domain is not expressed, this activity could relate to an active state of the phosphatase when SH2-mediated inhibition is already released. GRB2 weakly binds a longer CABIT2 containing the PRS. Reported pulldowns with full-length THEMIS showed robust association, suggesting that the CABIT1 domain also contributes to GRB2 binding.

The disease and therapeutic relevance of THEMIS remains understudied. THEMIS has been found to modulate the response to chimeric antigen receptor (CAR) T-cell therapy, making the study of the function of THEMIS also important to immuno-oncology.^38^ However, more structure-function studies are needed of other CABIT domains and the CABIT domain tandem to realize the therapeutic potential of THEMIS.

## Acknowledgements

T.S.B. was partially supported by a Ruth L. Kirschstein National Research Service Award (F32CA247198). T.S.B. acknowledges startup funds from Emory University School of Medicine and the Winship Cancer Institute of Emory University. This structural work was conducted at the Northeastern Collaborative Access Team beamlines (P30 GM124165, P41 GM103403) utilizing resources of the Advanced Photon Source at the Argonne National Laboratory (DE-AC02-06CH11357).

## Author Contributions

T.S.B, Z.Y, I.K.S., J.K.R., L.K.R, and D.O. purified proteins. T.S.B, Z.Y., and J.K.R. crystallized CABIT2. T.S.B. and M.J.E determined the CABIT2 structure. T.S.B., I.K.S., J.K.R., L.K.R performed phosphatase assays. D.O. performed pulldown experiments, L.K.R. performed DSF experiments. T.S.B, I.K.S., and M.J.E. wrote the manuscript. All authors have reviewed and approved the contents of this manuscript.

## Conflicts of Interest

The Eck lab receives (or has received within the past two years) sponsored research support from Novartis and Springworks Therapeutics.

## Data Availability

Coordinates and structure factors for the THEMIS CABIT2 domain will be deposited in the Protein Data Bank and publicly released upon final journal publication. Other data and materials are available upon reasonable request.

## Materials and Methods

### Protein expression and purification

Human THEMIS CABIT2 domain for crystallography (residues 267-550) with an N-terminal strep tag followed by a TEV protease cleavage site with the sequence MDWSHPQFEKSAVDENLYFQGG(THEMIS 267-550) was cloned into pEThisTT and verified by sequencing. THEMIS CABIT2 constructs spanning 267-550, 267-564, 267-530, 267-520, and 267-493 were cloned with an N-terminal His tag followed by a TEV protease cleavage site into pMCSG7 and verified by sequencing. THEMIS CABIT2 (residues 267-564) or SHP1 for pulldown experiments were cloned into pMCSG10 and purified with a N-terminal His and glutathione S-transferase (GST) tags. Full-length human SHP1 (PTPN6), SHP1 PTP domain (residues 243-530), and full-length human GRB2 were similarly cloned into pMCSG7 and verified by sequencing.

All proteins were expressed in *E. coli* Rosetta(DE3) cells at 18 ºC overnight after isopropyl β-d-1-thiogalactopyranoside (IPTG) induction. Cells were pelleted and resuspended in lysis buffer composed of 50 mM Tris pH 8.0, 200 mM NaCl, 1 mM tris(2-carboxyethyl)phosphine (TCEP), and 5% glycerol, lysed via sonication, and centrifuged at >50,000 g for 30 min. The resulting clarified supernatant was slowly flowed through a column containing Strep-Tactin MacroPrep resin for the strep-tagged construct or Ni-NTA resin for His-tagged constructs. The resin was washed with lysis buffer and eluted with lysis buffer containing 2.5 mM desthiobiotin or 200 mM imidazole for strep- and His-tagged proteins, respectively. Eluted protein was further purified via size exclusion chromatography on a prep-grade Superdex S200 column in 50 mM Tris pH 8.0, 200 mM NaCl, 1 mM TCEP, and 5% glycerol. Proteins were concentrated to 4-5 mg/mL by absorbance at 280 nm, flash frozen in liquid nitrogen, and stored at -80 ºC.

### Crystallization and structure determination

Crystals of THEMIS CABIT2 (267-550) with attached N-terminal strep tag and TEV protease cleavage site were obtained in numerous conditions from commercial screens. Crystals from D2 of the Classics Suite (Qiagen) were optimized via hanging drop vapor diffusion and large (>400 µm in length) hexagonal cylinder crystals were obtained in 0.1 M imidazole pH 6.9 and 1.0 M sodium acetate after 3-5 days with a protein concentration of 2-4 mg/mL. Derivatized crystals were soaked overnight in 0.1 M imidazole pH 6.9 and 1.0 M sodium acetate containing 1 mM potassium dicynaoaurate(I) (K[Au(CN)2]). Prior to freezing, crystals were cryoprotected in a solution of 0.1 M imidazole pH 6.9, 1.0 M sodium acetate, and 20% glycerol.

Diffraction data were collected at 100 K at the Advanced Photon Source at the Argonne National Laboratory on NE-CAT beamline 24-ID-C. Data for derivatized crystals were collected at 1.03987 Å, which is slightly above the wavelength corresponding to the L-III edge for gold anomalous signal. Data were indexed, integrated, scaled, and merged using Dials via xia2 compiled through SBGrid while keeping Bijvoet pairs separate to preserve anomalous signal.^39–41^ Experimental phasing was accomplished using AutoSol and initial models built using AutoBuild in Phenix.^42^ Refinements were performed using Phenix with iterative rounds of manual model building in Coot.^43^ For the native data set, models generated from experimental phasing were successfully used for molecular replacement (MR). The predicted CABIT2 domain structure from AlphaFold was also used to perform MR to compare with the model built from experimental phasing.^31,44^ Surface electrostatics were calculated using the APBS plugin^45,46^ for PyMol and conservation calculated using the ConSurf server.^34,35^

### Phosphatase assay

The activity of full-length SHP1 phosphatase or the protein tyrosine phosphatase (PTP) domain was measured via a fluorescence-based assay using 6,8-difluoro-4-methylumbelliferyl phosphate (DiFMUP). SHP1 phosphatase was used at a constant concentration of 0.5 or 3.5 µM for isolated PTP domain and full-length wild type, respectively. A range of THEMIS concentrations were prepared via serial dilution. All proteins and reagents were prepare in buffer composed of 50 mM Tris pH 8.0, 200 mM NaCl, 1 mM TCEP, and 5% glycerol. Six concentrations of THEMIS were used alongside THEMIS only and SHP only controls. Protein(s) were incubated in a volume of 10 µL at 2x final concentration in a white 384-well low-binding plate for 30 minutes at room temperature. Reactions were initiated by the addition of 10 µL DiFMUP at 10 µM for a final reaction volume of 20 µL and DiFMUP concentration of 5 µM. Data were immediately recorded every minute on a CLARIOstar Plus (BMG Labtech) with an excitation wavelength of 358 nm and emission wavelength of 620 nm for 30 min as the reaction progressed.

### Thermal shift assay

DSF was performed using His-tagged CABIT2 domain (267-564) at a concentration of 5 µM and SYPRO Orange dye at 2.5x concentration (from 1000x stock, Sigma-Aldrich) in buffer containing 50 mM Tris pH 8.0, 200 mM NaCl, 5% glycerol, 1 mM TCEP, and 1% DMSO and a final volume of 10 µL. Samples were heated from 35-65 °C at a rate of 2 °C/min in an Eppendorf realplex 4 Mastercycler S qPCR instrument and fluorescence detected with excitation and emission wavelengths of 552 and 578 nm, respectively. The melting point (T_m_) corresponded to the inflection point of the resulting sigmoidal melt curve.

### Pulldowns

Pulldowns were performed using GST-tagged CABIT2 domain and His-tagged GRB2 or SHP1. GST-CABIT2 or GST-SHP1 were immobilized on glutathione-agarose resin and incubated with equimolar amounts of His-tagged GRB2 or CABIT2, respectively. Resin was washed with buffer containing 50 mM Tris pH 8.0, 200 mM NaCl, 5% glycerol, and 1 mM TCEP and eluted with buffer supplemented with 10 mM reduced glutathione. Proteins were separated on a 12% SDS-PAGE and visualized by Coomassie brilliant blue staining.

## Notes

### Competing Interest Statement

The authors have declared no competing interest.

