## Supplemental Information for "Structure of the CABIT2 domain of THEMIS reveals a novel protein fold with an inserted SH3-like domain"

**SI Table 1:** Crystallographic data collection and refinement statistics.

| **Data Collection** | |
|---|---|
| **Wavelength (Å)** | 1.03987 |
| **Number of merged data sets (single crystal)** | 3 |
| **Resolution range (Å)** | 62.45 - 2.90 (3.00 - 2.9) |
| **Space group** | $P6_122$ |
| **Unit cell** | |
| **a, b, c (Å)** | 72.96 72.96 409.87 |
| $\alpha, \beta, \gamma$ **(°)** | 90 90 120 |
| **Total reflections** | 2331826 (235270) |
| **Unique reflections** | 15442 (1482) |
| **Multiplicity** | 151.0 (158.8) |
| **Completeness (%)** | 97.66 (88.19) |
| **Mean I/σ(I)** | 60.59 (1.47) |
| **Wilson B-factor** | 117.22 |
| $R_{merge}$ | 0.2929 (4.097) |
| $R_{pim}$ | 0.0255 (0.322) |
| $CC_{1/2}$ | 0.967 (0.81) |
| **Anomalous completeness (%)** | 100.0 (100.0) |
| **Anomalous multiplicity** | 84.7 (86.0) |
| $CC_{anom}$ | 0.942 (0.016) |
| **Anomalous slope** | 1.368 |
| **Refinement** | |
| **Reflections used in refinement** | 14772 (1221) |
| **Reflections used for** $R_{free}$ | 1467 (122) |
| $R_{work}$ | 0.245 (0.626) |
| $R_{free}$ | 0.268 (0.664) |
| **Number of non-hydrogen atoms** | 2329 |
| **macromolecules** | 2329 |
| **Protein residues** | 289 |
| **RMS bonds (Å)** | 0.002 |
| **RMS angles (Å)** | 0.55 |
| **Ramachandran favored (%)** | 91.64 |
| **Ramachandran allowed (%)** | 6.62 |
| **Ramachandran outliers (%)** | 1.74 |
| **Rotamer outliers (%)** | 2.27 |
| **Average B-factor** | 123.46 |

Statistics for the highest-resolution shell are shown in parentheses.

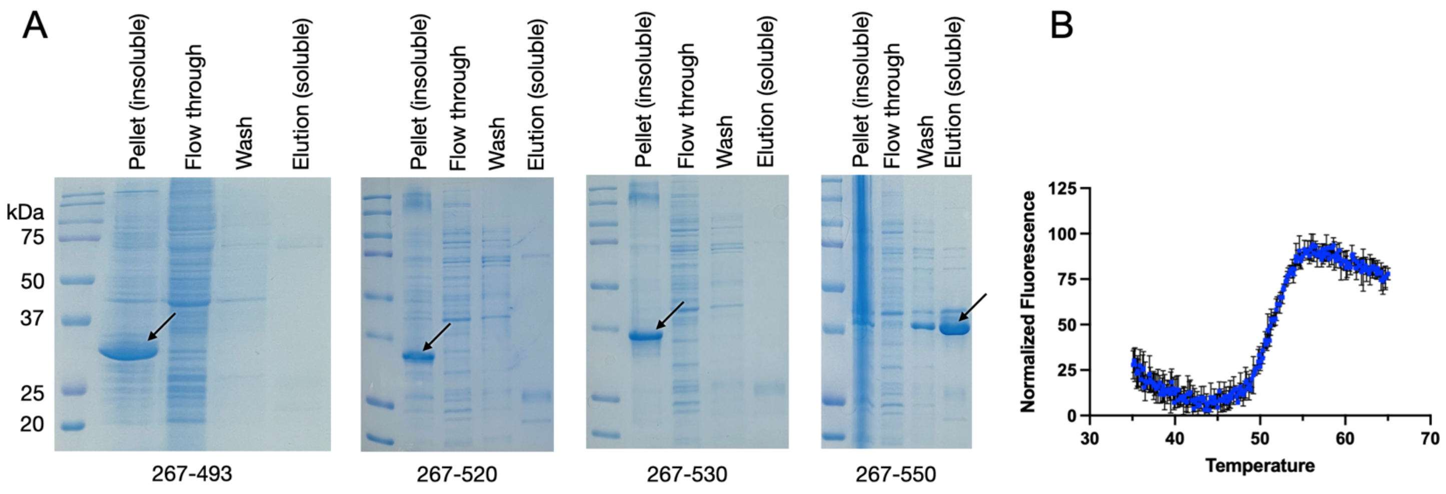


**SI Figure 1:** Purification and thermal stability of CABIT2 constructs. A) Purification of 6xHis-tagged CABIT2 constructs with varying C-terminal cutoffs. Only 267-550 results in soluble protein that can be eluted from Ni resin. B) The CABIT2 construct crystallized (residues 267-550) has a melting temperature of $T_m$=51.34 ± 0.03 °C as determined by differential scanning fluorimetry (DSF) (n=3, mean±SD).

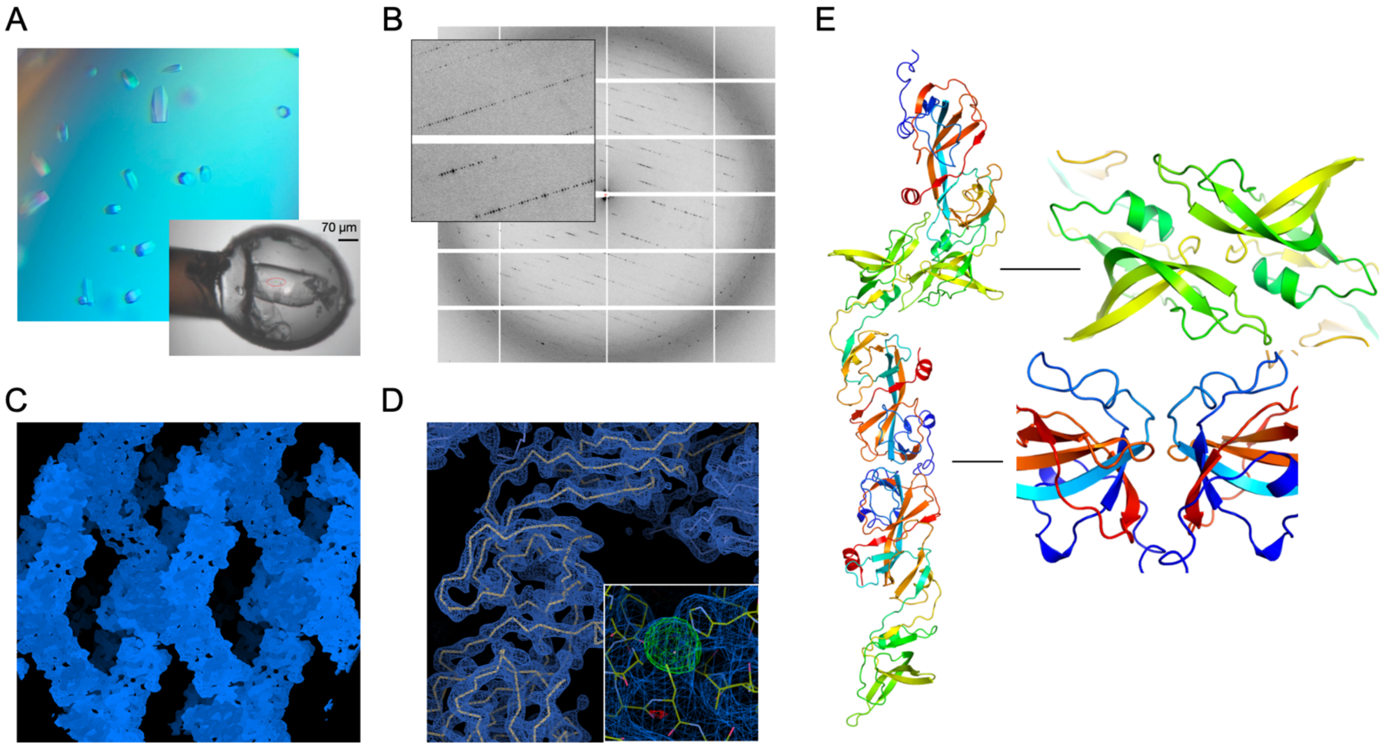


**SI Figure 2:** Details of CABIT2 domain crystallization and structure determination. A) CABIT2 crystals grow with a hexagonal cylinder morphology (0.1 M imidazole, pH 6.5-7.0, 0.8-1.2 M sodium acetate. Inset is a picture of the gold-soaked crystal that was used to determine the structure. B) Diffraction pattern from gold-soaked crystal. The large unit cell axis produces close spots that are highlighted in the inset photo. C) Lattice density ($2F_o$-$F_c$ at 1 $\sigma$) following SAD phasing and statistical density modification. D) Backbone building into density ($2F_o$-$F_c$ at 2 $\sigma$). Inset image shows strong positive $F_o$-$F_c$ density for a gold ion upon structure refinement. E) Portion of the crystal lattice showing crystal contact details. Head-head and core-core interfaces primarily drive lattice formation.

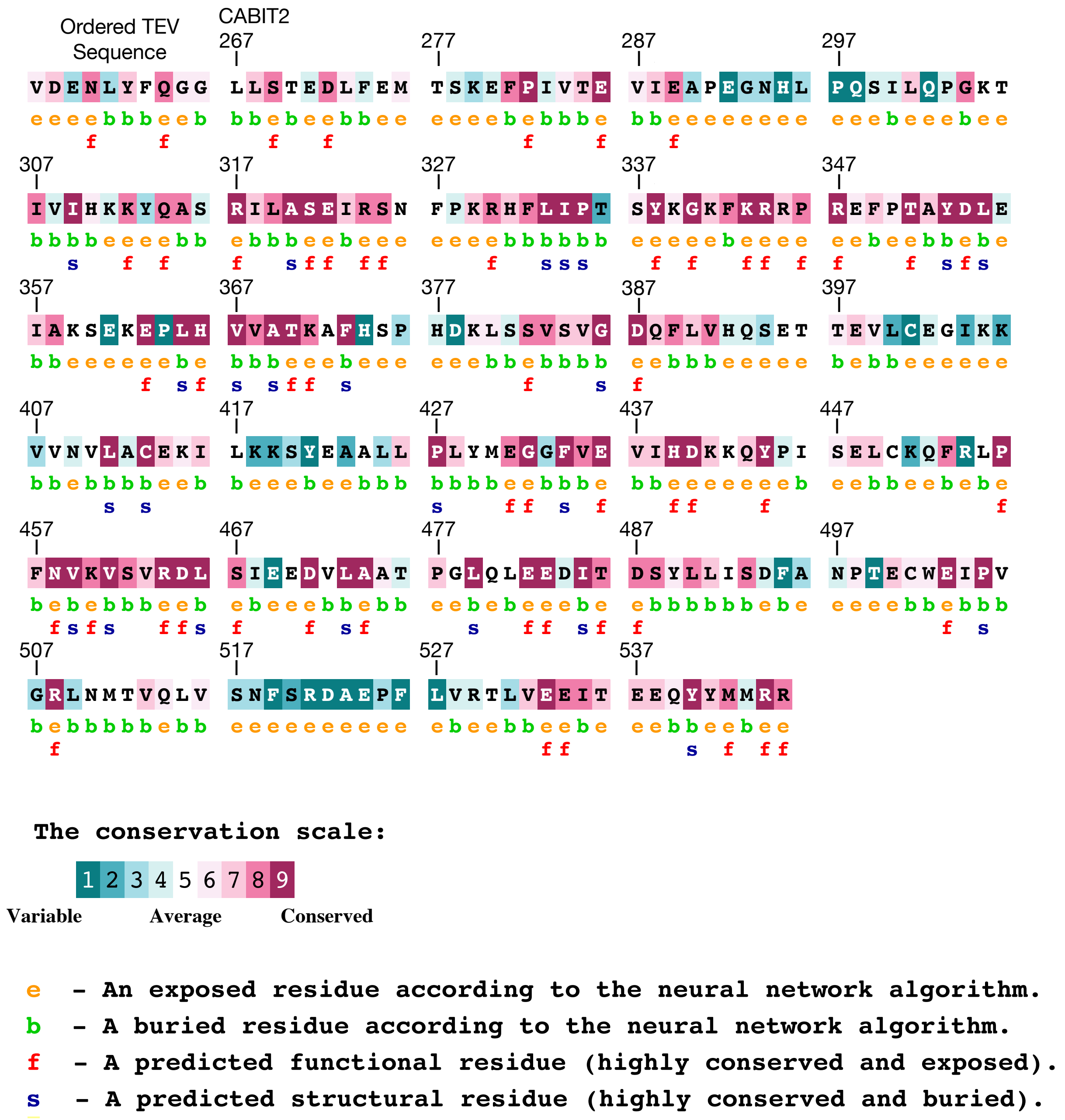


**SI Figure 3:** Conservation scores from the CONSURF server mapped onto the primary sequence of the resolved residues in the crystallized CABIT2 domain.

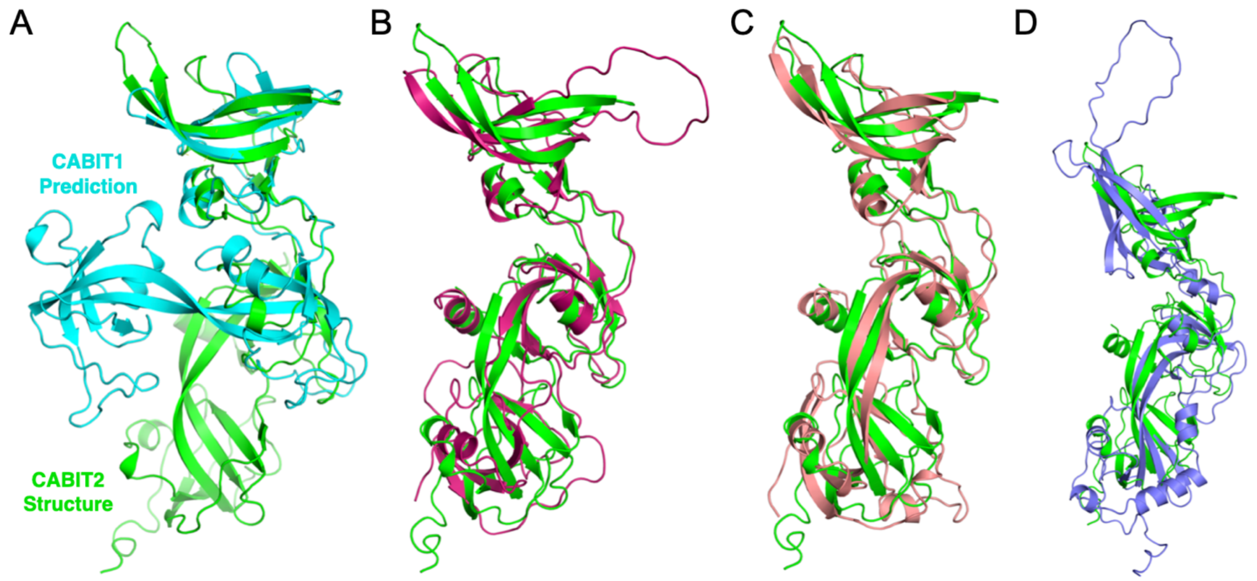


**SI Figure 4:** Comparison of THEMIS CABIT2 with other CABIT isoforms. A) Structural superposition between the AlphaFold 3 predicted THEMIS CABIT1 domain and experimental CABIT2 structure. Domains are aligned to the SH3-like head subdomain. B) Superposition with predicted THEMIS2 CABIT2 (magenta, RMSD=3.3 Å). C) Superposition with predicted mouse THEMIS3 CABIT2 (salmon, RMSD=3.6 Å). THEMIS3 is not expressed in primates. D) Structural superposition with GAREM CABIT domain (purple, RMSD=10.7 Å).
